# Quantifying the effects of recent glacial history and future climate change on a unique population of mountain goats

**DOI:** 10.1101/2022.01.19.476931

**Authors:** Kiana B. Young, Tania M. Lewis, Kevin S. White, Aaron B.A. Shafer

## Abstract

Human disturbance and climate change can impact populations by disrupting movement corridors and reducing important habitat. Characterizing how animals respond to such environmental changes is valuable for conservation as many species, especially habitat specialists, can experience reduced genetic diversity when deleterious habitat change occurs, leading to an increased likelihood of extirpation. Mountain goats (*Oreamnos americanus*) exemplify this conservation challenge; their geographically isolated habitat can inhibit gene flow, making them susceptible to population declines in the face of anthropogenic-induced landscape change. To facilitate biologically informed population management of mountain goats in Glacier Bay National Park, Alaska, we characterized the fine-scale genetic population structure and examined how future climate change could impact the population density of these mountain goats. We used DNA samples to estimate diversity and depict the genealogical history. Climate response models allowed us to simulate changes to suitable habitat and predict how this might influence future population structure. Our results indicated that three genetically distinct subpopulations exist in Glacier Bay and that the population structure is reflective of the historic landscape patterns. Climate modeling predicted that demographic productivity was likely to be reduced for all subpopulations; additionally, we found that climate change likely degrades the suitability of movement corridors that facilitate gene flow between subpopulations, ultimately increasing the cost of travel. Understanding such fine-scale patterns are key to managing subpopulations, particularly with impending changes to the landscape.

## 1. INTRODUCTION

The colonization of landscapes is a central question in biogeography (Lomolino et al. 2017) that can often be revealed by mapping the genetic diversity and differentiation of populations. For example, plotting genetic diversity across the landscape can retrace the route individuals took (i.e. stepping-stone model; Kimura and Weiss 1964, Baltazar-Soares et al. 2020), while more complex models can estimate divergence times and infer population size changes (i.e. Csilléry et al. 2010). When colonization follows a disturbance, quantifying the genetic response can aid in predicting changes to the demography of that population as the landscape continues to change. This information can help inform management decisions, for example, by minimizing disturbance to those subpopulations with lower genetic diversity (Bouzat 2010, Stronen et al. 2019).

Landscape disturbances come in a variety of forms, and while each type of disturbance has unique characteristics, there are similarities between regimes (Newman 2019). Glaciation and subsequent deglaciation events provide opportunities to learn about animal movement across broad scales (Hewitt 1999). The most recent large-scale ice age, during the Pleistocene Epoch, ended around 11,700 years ago, and led to a range of species’ responses, particularly in North America (Lister 2004, Pearson 2013, Bibi and Kiessling 2015, Puzachenko and Markova 2019). During the Last Glacier Maximum, two major refugia existed in western North America: Beringia and the Pacific Northwest (Hultén 1937, Pielou 1991). While those areas are widely accepted as refugia for species during this ice age, there is evidence of additional, smaller refugia along coastal Alaska and British Columbia (Shafer et al. 2010). Beyond the major ice ages, smaller-scale glaciation events have also occurred in North America. Notably, Glacier Bay National Park and Preserve (GBNPP), located in Southeast Alaska, experienced a small glaciation event, a product of the Little Ice Age, only ∼300 years ago (Connor et al. 2009). The native wildlife was forced to leave the area that is now Glacier Bay fjord to escape the advancing glacier. Glaciers have since retreated, allowing flora and fauna to recolonize the landscape (Milner et al. 2007, Boggs et al. 2010).

The origin of many recolonized species in GBNPP is unknown. Brown bears (*Ursus arctos*) and black bears (*Ursus americanus*) recolonized GBNPP from both the northeast and northwest after the Little Ice Age with Glacier Bay fjord and glacier-covered mountains acting as barriers to dispersal (Lewis et al. 2015, 2020). While other mammals might have recolonized in a similar way, the habitat associations of a species likely play a role in their movement across the landscape. Bears, for example, might travel along coastlines or through forests, while alpine ungulates such as mountain goats (*Oreamnos americanus*) likely select high elevation, mountainous terrain leading to different movement patterns (Rice 2008). However, marine waterways and steep glacier covered mountains or icefields likely inhibit movement of mountain goats. Interestingly, while mountain goat habitat selection models suggest avoidance of glaciers in coastal Alaska (Shafer et al. 2012), both expert opinion and landscape modeling suggested a minimal effect of glaciers on movement in the Cascade Mountains (Shirk et al. 2010).

Reconstructing the historical colonization of a species can help predict future movements in response to disturbance and is particularly relevant as land management agencies develop plans to mitigate impacts to vulnerable and climate sensitive wildlife populations. Climate change is expected to result in widespread changes in glaciers and landscape configurations and learning about those processes in a place like GBNPP, where major deglaciation has occurred in recent times, provides an important opportunity to gain insights about how predicted climate change may impact ecological systems more broadly. Here, we used non-invasive genetic samples to examine the population genetic structure of mountain goats in and surrounding GBNPP. We use a Bayesian computational approach to reconstruct the demographic history and climate projections to model how these subpopulations and movement corridors might be impacted by climate change.

## 2. METHODS

### 2.1 Study area and sample collection

Glacier Bay National Park and Preserve is located in Southeast Alaska and is characterized by glacial- and river-made valleys surrounded by mountain peaks and fjords (Boggs et al. 2010). Within the boundary of GBNPP, four mountain goat study areas were identified based on geographic patterns of distribution, abundance, and management interest. These study areas were Mount Wright, Tidal Inlet, Marble Mountain, and Table Mountain (Figure 1). We included samples from the adjacent Haines-Skagway and the Bering Glacier areas (Figure 1). These latter two areas were studied because they were considered putative source populations for Glacier Bay.

**Figure 1.**
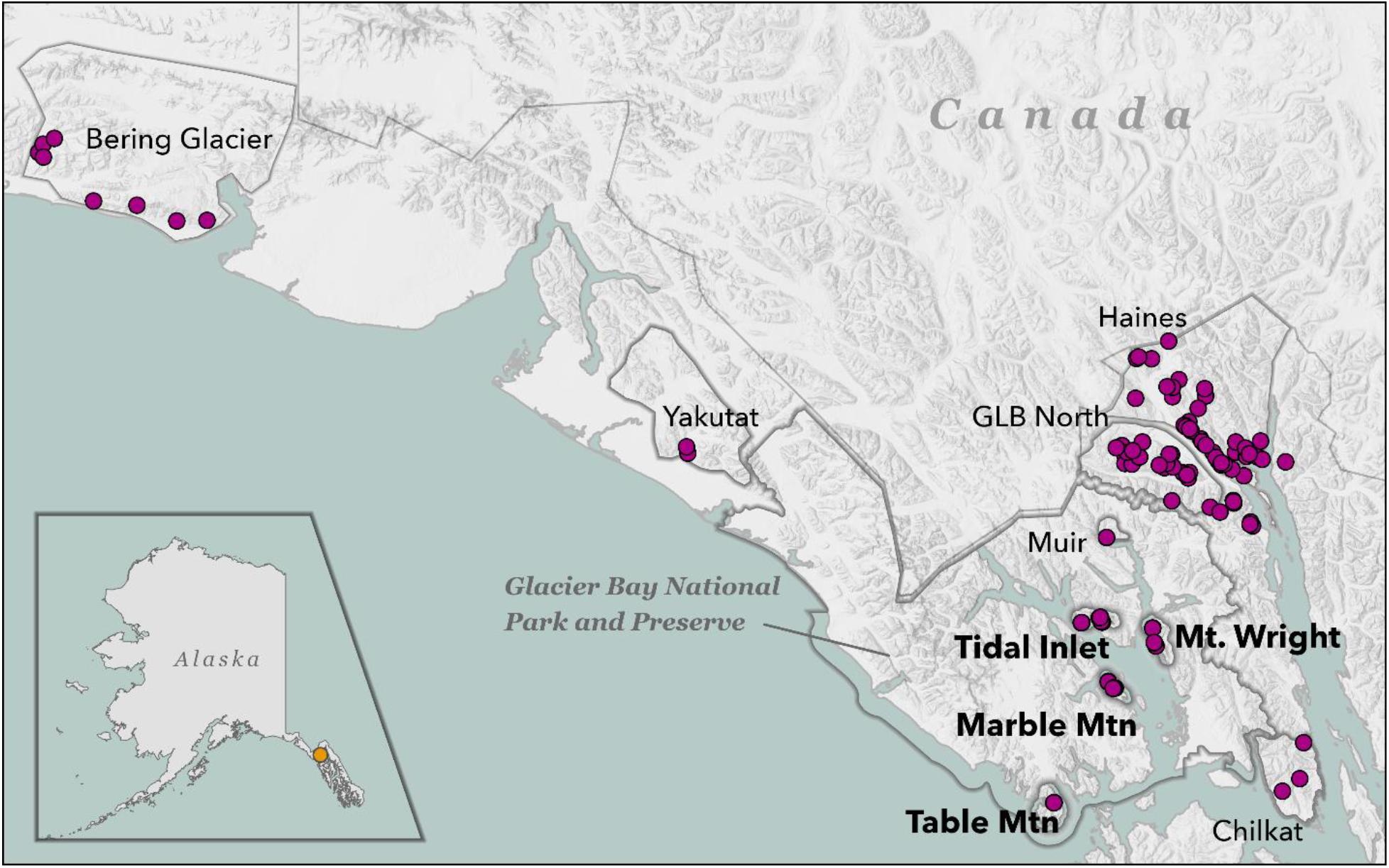
Map of mountain goat (*Oreamnos amercanus*) samples for Glacier Bay National Park, Alaska analysis. The four focal study areas are indicated in bold. All other study areas (not in bold) were used for demographic and admixture analysis.

During the summers of 2017 through 2020, we collected pellet samples from GBNPP. Once a pellet group was determined to be fresh using protocols from Poole et al. (2011), we swabbed the outside of multiple pellets from the same pellet group and stored the swabs in a vial of Longmire’s solution, a lysis buffer designed to preserve DNA and prevent contamination (100mM Tris, 100mM EDTA, 10mM NaCl, 0.5% SDS, 0.2% sodium azide). Samples were stored at -20 degrees C. Tissue samples adjacent to GBNPP were collected by Alaska Department of Fish & Game (ADFG) through harvest and research sampling following field sampling methods described in Shafer et al. (2011) and White et al. (2021a, 2021b).

### 2.2 DNA extraction and lab methods

DNA was extracted using a Qiagen DNeasy Blood and Tissue Kit (Qiagen Inc., Valencia, California, USA), following the manufacturer’s protocols. We amplified 17 polymorphic microsatellites over 3 multiplex PCR pools using previously published non-invasive genotyping protocol (White et al. 2021a). We used the software program Geneious (v 10.1.2, Kearse et al. 2012) to manually call alleles requiring a minimum strength of 250 RFUs. Pellet samples were genotyped in triplicate and samples where fewer than two of the replicate genotypes matched were dropped following the approach of White et al. (2021a). Samples with genotypes less than 80% complete were dropped from the analysis. Positive and negative controls were included in all steps.

### 2.3 Genetic variation

We used the R package ALLELEMATCH (Galpern et al. 2012) to determine if individuals were sampled multiple times (alleleMismatch = 2). To test for deviations from Hardy-Weinberg Equilibrium (HWE) and linkage disequilibrium (LD), we used the R package genepop (Rousset 2008). We checked for allelic dropout using the program Micro-Checker (v2.2.3 Van Oosterhout et al. 2004). For each study area we estimated the mean and private allelic richness using ADZE.

GENHET (Coulon 2010) and adegenet (Jombart 2008) were used to estimate individual- and subpopulation-level heterozygosity, respectively. We estimated the effective population size (*N*_*e*_) using NeEstimator (v2.1, Do et al. 2014), and used the program SPAGeDi (Hardy and Vekemans 2002) to calculate the *F*_*ST*_ and Nei’s *D*. We calculated spatial autocorrelation using the parameter Moran’s *I* at 5 km distance bins.

### 2.4 Population structure and demographic history

We visualized clusters of samples using a principal components analysis (PCA) and plotted individual principal component scores against latitude and longitude as a proxy for isolation-by-distance. The Bayesian clustering program STRUCTURE (Pritchard et al. 2000, Falush et al. 2003, 2007, Hubisz et al. 2009) was used to identify subpopulations using the admixture model with independent runs from K=1 to K=20 and a burn-in period of 5 × 10^5^ and 1 × 10^6^ MCMC iterations. We used a combination of Evanno et al. (2005) and Puechmaille (2016) methods determine the optimal K. Admixed individuals were considered when q < 0.8.

The Bayesian approximate computation software DIYABC (v2.0, Cornuet et al. 2014) was run to reconstruct the demographic history both within GBNPP and GBNPP and adjacent populations (Bering Glacier and Haines-Skagway) where we assessed four different recolonization scenarios (Figure S1, S2). We simulated 1 × 10^5^ datasets for each scenario and recorded the following summary statistics: mean number of alleles, genic diversity, mean Garza-Williamson’s M, and *F*_*ST*_. Parameters were adjusted based on their posterior distribution and the optimal model was selected by using measures of posterior probabilities of scenarios. Time estimates in number of generations were converted to years by multiplying by a generation time of 6 years following Martchenko et al. (2020).

### 2.5 Climate change predictions

We used a population model developed by White et al. (2018) to simulate demographic responses of each genetically distinct subpopulation to predicted changes in climate. This approach simulates climate effects by modeling how changes in mean July/August temperature and total annual snowfall [derived from precipitation as snow (PAS)], influence sex- and age-specific survival (White et al. 2011); a key predictor of population performance (Hamel et al. 2006). Temperature and PAS predictions were generated with the ClimateNA software package (v5.10, Wang et al. 2016), and baseline values were calculated by taking the average temperature and PAS over the past 20 years. Five GCM models were used to predict the changes in temperature and PAS: CCSM4, GFDL-CM3, GISS-E2-H, IPSL-CM5B-LR, and MRI-CGCM3 (White et al. 2018). For each model we considered two emission scenarios (RCP 4.5 and RCP 8.5), and predicted out to year 2025, 2055, and 2085. Snowfall variability was modeled using total annual snowfall measurements collected at the Gustavus Airport during 1965-2019. Subpopulation size for each study area was derived based on aerial survey data collected during 2012 (Lewis and White 2015) and adjusted for sightability bias following analytical methods described in White et al. (2016). The stable age distribution was used to determine the age and sex structure for each initial population size following White et al. (2018). Age-specific density dependent fecundity and kid survival was also parameterized as per White et al. (2018). We ran each model 1000 times and calculated how many times the subpopulation size fell below *N* = 2 by the year 2085 to estimate the probability of quasi-extinction, a general metric of population performance over time.

We used ecological niche modelling (ENM) and Least Cost Path (LCP) analysis to predict how movement corridors might change in response to climate change. We used coordinates of both the invasive and non-invasive research samples collected in the field combined with a suite of climate raster data layers. The current and future cost landscape rasters were created using the ENM software Maxent (v3.3.3, Phillips et al. 2006) and ArcGIS Pro (v 2.6.3) software. We evaluated the predictive capability of the model using cross-validation methods with 75% of the samples as training samples to build the model; the remaining 25% of the samples were omitted from the model building process and later used to test the model. We ran the model using a variety of combinations of parameters and compared Area Under the Curve (AUC) to measure model performance. We used 19 bioclimatic variables, downloaded from WorldClim (https://www.worldclim.org/) for current climate data. Additionally, we used landcover type and digital elevation model (DEM) rasters downloaded from the United States Geologic Survey database (https://apps.nationalmap.gov/downloader/#/). The DEM raster was used to calculate the heat load index which is a measure of the incident radiation in a location based on the slope and aspect; this attribute has previously been found to affect mountain goat space use (Shafer et al. 2012) and is subject to change with climate. All layers had a resolution of 30 arc-seconds. A jacknife test of variable contributions was used to determine which variables contributed to the model; variables that did not contribute were removed and the model was re-run. Future climate variables were downloaded from the GCM Downscaled Data Portal (http://www.ccafs-climate.org/data_spatial_downscaling/). We used the Representative Concentration Pathways models RCP 4.5 and RCP 8.5 from the GFDL_CM3 model projected for the year 2080 (Donner et al. 2011), used in the IPCC Fifth Assessment Report (Shukla et al. 2019).

To calculate the LCP between the locations of each research sample collected in northern Southeast Alaska, we used the R package ‘gdistance’(van Etten 2017). Least cost path matrices were produced and compared for current, future (year 2085) under the RCP 4.5 scenario, and future (year 2085) under the RCP 8.5 scenario. We compared the cost distance to the genetic distance parameter Moran’s *I* to determine how the genetic distance changed as the spatial LCP distance increased and the cost distance to the Euclidean distance to compare the cost and distance values. We used a multiple regression on distance matrices (MRM) method in the R package ‘ecodist’ (Goslee and Urban 2007) that used Euclidean distance and current LCP distance as predictor variables, with the genetic distance as the response variable.

## 3. RESULTS

### 3.1 Population genetic diversity statistics

A total of 68 unique samples from pellets in all four sampling areas in GBNPP were genotyped at 17 polymorphic microsatellites (Table 1). An additional 69 (3 pellet, 66 tissue) samples were added from areas surrounding the GBNPP focal areas to investigate connectivity of mountain goats across the GBNPP boundary. Nei’s *D* between the sampling areas ranged from 0.01 to 0.18 with the lowest levels of genetic differentiation between Mt. Wright and Tidal Inlet and the highest between Marble Mountain and Table Mountain (Table 2). The PCA showed clustering of individuals according to sampling area, but Mt. Wright and Table Mountain samples overlapped (Figure 2A). The first principal component versus latitude and longitude analysis did not show a clear relationship (p > 0.05, Figure S3); however, an IBD and spatial autocorrelation pattern was present, with genetic relatedness decreasing slightly as spatial distance increased (Figures S4, S5). There was no correlation between *H*_*O*_, *F*_*IS*_, and *N*_*e*_ and longitude or latitude in GBNPP (Figure S6).

**Table 1.** Diversity statistics for mountain goats in four sampling areas in Glacier Bay National Park, Alaska (n = 68). Data were generated using Genalex v6.503 and NeEstimator v2.1.

| *Sampling area* | *No. of samples* | *Assigned pop* | *Mean q-value* | *Observed heterozygosity* | *Expected heterozygosity* | *F<sub>IS</sub>* | *N<sub>e</sub> (95% CI)* |
| --- | --- | --- | --- | --- | --- | --- | --- |
| Marble Mountain | 25 | Marble Mtn | 0.87 | 0.19 ± 0.049 | 0.24 ± 0.056 | 0.18 ± 0.078 | 6.6-50.7 |
| Mt Wright | 10 | TidalInlet/<br>MtWright | 0.76 | 0.34 ± 0.050 | 0.36 ± 0.054 | 0.02 ± 0.044 | 8.3-∞ |
| Table Mountain | 14 | Table Mtn | 0.92 | 0.32 ± 0.069 | 0.34 ± 0.047 | 0.07 ± 0.126 | 2.5-∞ |
| Tidal Inlet | 19 | TidalInlet/<br>MtWright | 0.82 | 0.30 ± 0.052 | 0.33 ± 0.045 | 0.17 ± 0.095 | 5.5-21.5 |

**Table 2.**
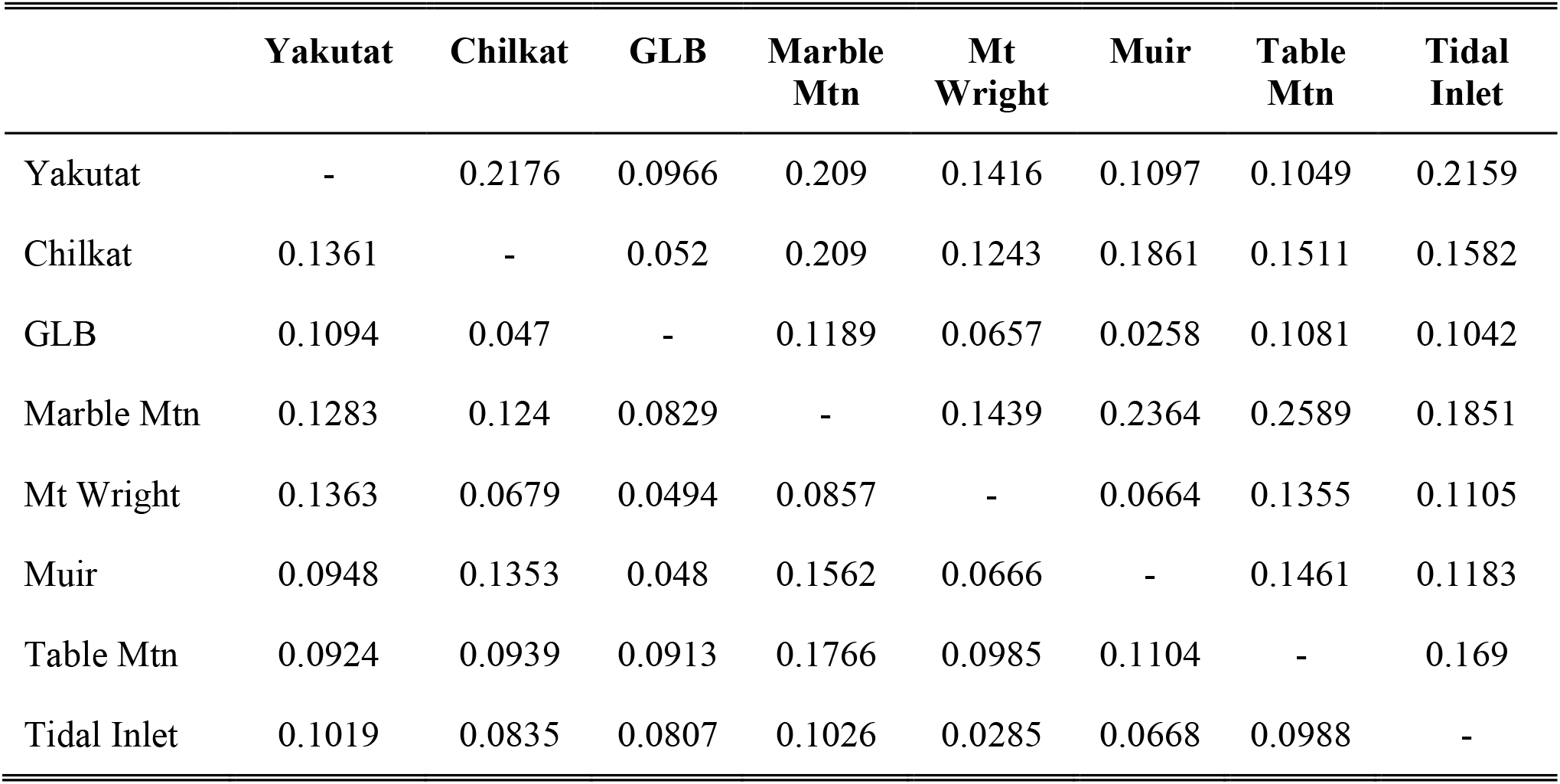
Pairwise Nei’s *D* (lower matrix) and *F*_*ST*_ (upper matrix) for mountain goats in eight sampling areas in and around Glacier Bay National Park, Alaska and (n = 137). Data were generated using SPAGeDi v1.5d.

**Figure 2.**
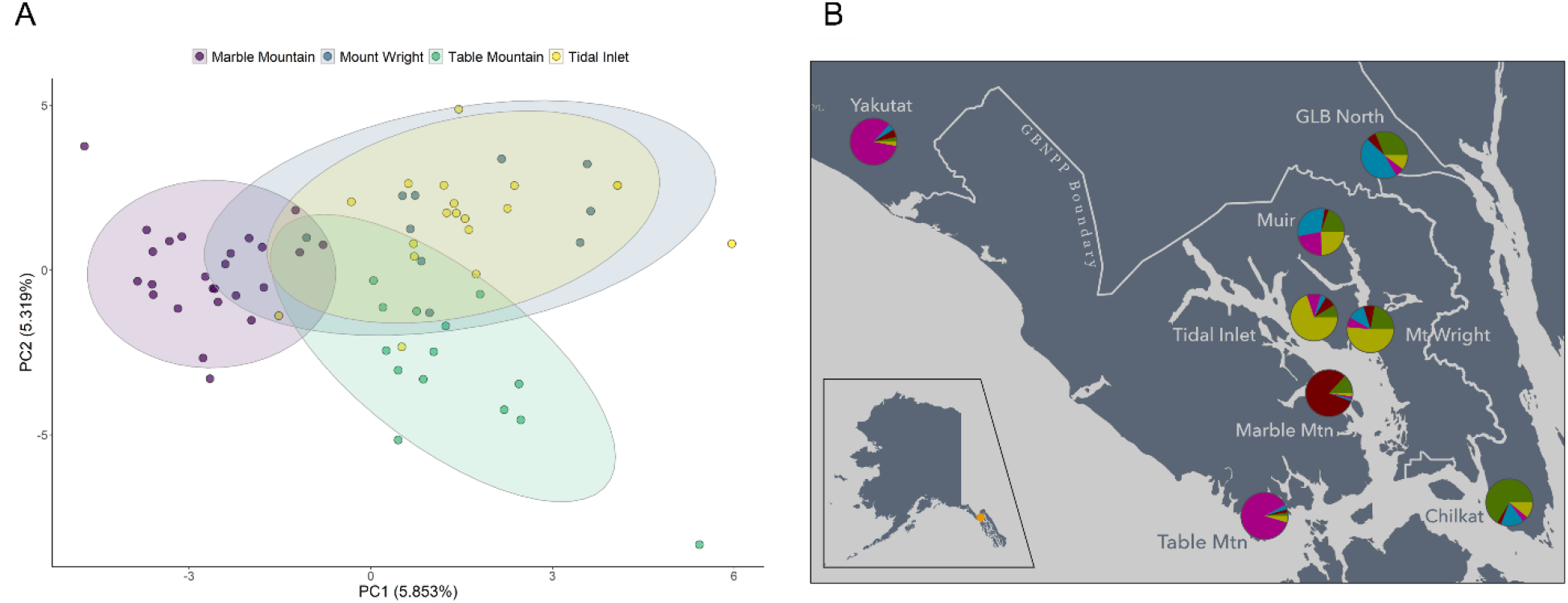
The population structure of mountain goats in and around Glacier Bay National Park and Preserve. A) Principal Components Analysis (PCA) of mountain goats from four sampling areas in Glacier Bay National Park, Alaska (n = 68). The proportion of variance explained by each axis is shown in parentheses. Subpopulations are delineated by color and ellipses. B) Average STRUCTURE subpopulation assignments for mountain goats in each of the four study areas in Glacier Bay National Park, Alaska and areas surrounding Glacier Bay. Pie charts indicate the q-value assignments for each K=5. The four GBNPP sampling areas are indicated in bold.

### 3.2 Population structure and demography

STRUCTURE analysis supported three distinct subpopulations of mountain goats in GBNPP (Figure 2B, S7). When surrounding individuals were included in the analysis 5 subpopulations were detected, the same three within the GBNPP boundary and two intermixed subpopulations northeast of GBNPP (Figure 2B, S8). The amount of admixture was lowest for the three GBNPP subpopulations (proportion of individuals that were admixed: 0.16-0.33) and highest for the subpopulations to the north (0.38-0.41, Table S1). Demographic modelling indicated that within GBNPP, the Table Mountain subpopulation split off first ∼7,920 years ago (95% CI: 3,102-37,500). Following that split, the Tidal Inlet/Mt. Wright subpopulations split from Marble Mountain ∼1,050 years ago (95% CI: 524-7,080; Figure S1 Scenario 2). Results from the broad scale demographic analysis indicated that GBNPP mountain goats split off most recently from the Northeast in the Haines-Skagway area ∼12,480 years ago (95% CI: 5,460-45,600; Figure S2 Scenario 1).

### 3.4 Climate change population modeling

Subpopulations showed numeric declines under all climate scenarios. For all four sampling areas, the CCS and GFDL GCMs under the RCP 4.5 emissions scenarios showed subpopulations reaching quasi-extinction by the year 2085 (Table 3). The CCS, GDFL, and ISPL GCMs showed subpopulations reaching quasi-extinction for all runs in the RCP 8.5 scenario (Table 3). The optimal Maxent model (AUC = 0.942) was based off 118 sample locations and included 13 environmental variables (Table S2) and indicated that the ecological niche for mountain goats in Southeast Alaska, given their current location and these environmental layers, is reduced in response to climate change (Figure 3A). The mean cost of travel between all individuals for the current LCP analysis was 1.84 × 10^6^; conversely, mean cost of travel between all individuals for the future RCP 8.5 LCP analysis was 1.40 × 10^16^ (Figure 3B). Multiple regressions indicated that geographic distance is negatively correlated with Moran’s *I* (R^2^ = 0.22, p < 0.01), while including LCP did not improve the model fit (Table S3).

**Table 3.** Projected mountain goat population response to climate change over a 65-year period (2020-2085) for four subpopulations in Glacier Bay National Park, Alaska (total n = 68, Marble Mountain = 49, Mount Wright = 244, Table Mountain = 208, Tidal Inlet = 214). Subpopulations were simulated 1000 times under five global climate models (GCM) each with two emission scenarios (RCP 4.5 and 8.5). Initial population sizes were determined based on aerial surveys conducted in 2012 and corrected for sightability. Subpopulations were simulated 1,000 times each. *N* indicates the estimated population size of each study area.

| GCM | % quasi-extinction (N<2) by 2085 |  |  |  | Years to extinction |  |  |  |
| --- | --- | --- | --- | --- | --- | --- | --- | --- |
|  | *Marble Mtn* | *Mt. Wright* | *Table Mtn* | *Tidal Inlet* | *Marble Mtn* | *Mt. Wright* | *Table Mtn* | *Tidal Inlet* |
| **RCP 4.5** |  |  |  |  |  |  |  |  |
| **CCS** | 98.6 | 62.2 | 37.6 | 42.7 | >65 | >65 | >65 | >65 |
| **GFDL** | 100 | 88.4 | 83.0 | 62.5 | 55.51±0.15 | >65 | >65 | >65 |
| **GISS** | 0.0 | 0.0 | 0.0 | 0.0 | >65 | >65 | >65 | >65 |
| **ISPL** | 0.0 | 0.0 | 0.0 | 0.0 | >65 | >65 | >65 | >65 |
| **MRI** | 0.0 | 0.0 | 0.0 | 0.0 | >65 | >65 | >65 | >65 |
| **RCP 8.5** |  |  |  |  |  |  |  |  |
| **CCS** | 100 | 100 | 100 | 100 | 45.02±0.09 | 47.97±0.09 | 49.93±0.10 | 50.59±0.09 |
| **GFDL** | 100 | 100 | 100 | 100 | 48.10±0.10 | 52.12±0.10 | 54.17±0.10 | 52.00±0.10 |
| **GISS** | 0.0 | 0.0 | 0.0 | 0.0 | >65 | >65 | >65 | >65 |
| **ISPL** | 100 | 100 | 100 | 100 | 52.14±0.09 | 56.44±0.08 | 57.65±0.09 | 57.60±0.09 |
| **MRI** | 0.0 | 0.0 | 0.0 | 0.0 | >65 | >65 | >65 | >65 |
| **N** | 49 | 244 | 208 | 214 |  |  |  |  |

**Figure 3.**
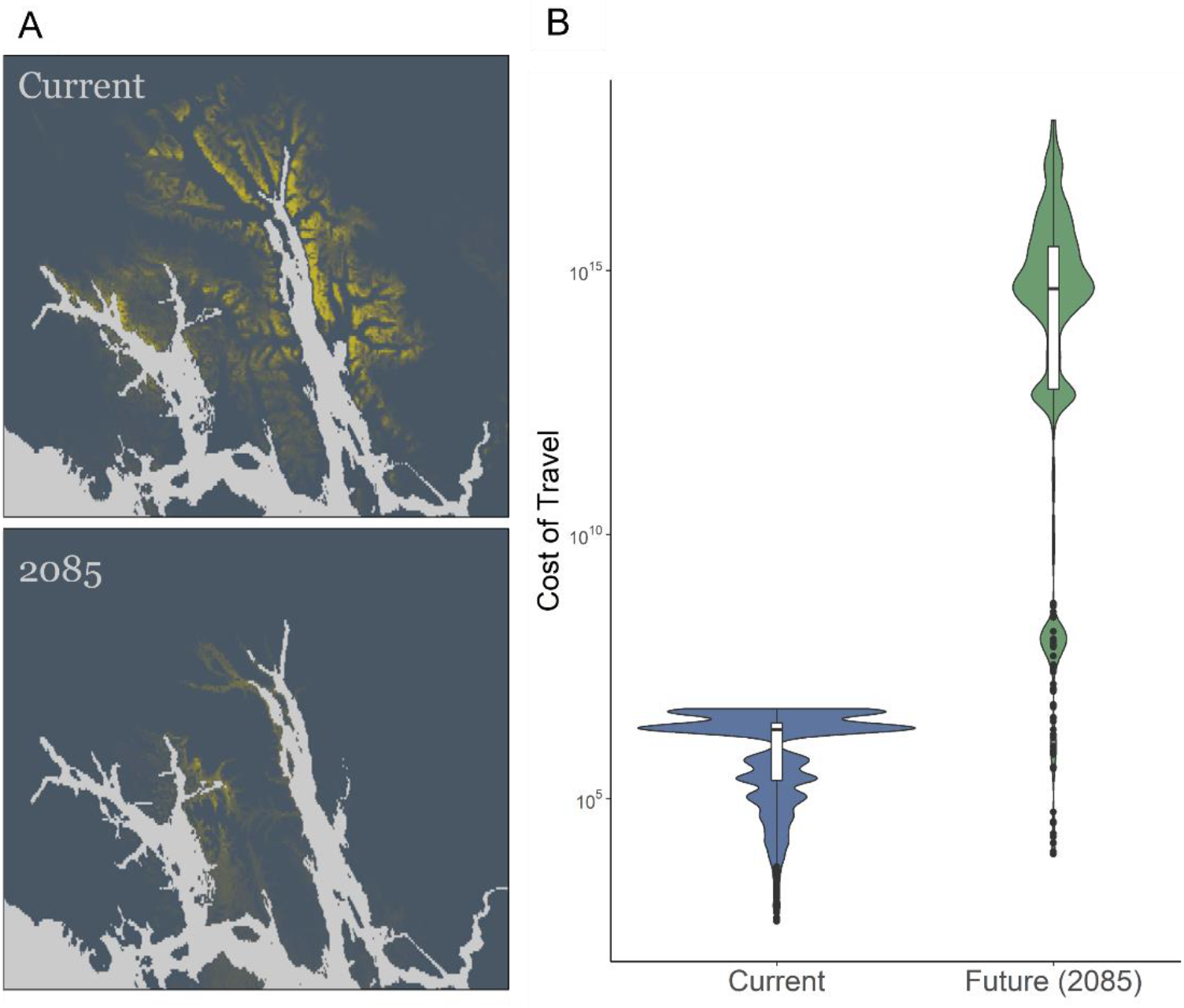
The effects of climate change on mountain goats in northern Southeast Alaska. (A) Maps of current (top) and year 2085 (bottom) habitat suitability for mountain goats in northern Southeast Alaska based off Maxent analysis. Yellow indicates more suitable habitat and blue indicates less suitable habitat. The global climate model GFDL-CM3 was used for this model. B) Violin plot of the average current and future cost of travel for mountain goats between each sample location in northern Southeast Alaska. Costs were calculated using Maxent suitability layers and Least Cost Path analysis.

## 4. DISCUSSION

### 4.1 Patterns of genetic differentiation

Alaska has a dynamic geologic history with multiple glaciation events that have shaped the current landscape and biodiversity (Svenning et al. 2015, Antonelli et al. 2018). Our results suggest that the mountain goats in GBNPP originated from a local source population in Southeast Alaska rather than dispersing from Southcentral Alaska, or more distant refugial populations. Shafer et al. (2011) provided evidence for a hot spot of mountain goat genetic diversity around Haines, Alaska with peripheral populations appearing to radiate to the surrounding areas. We hypothesize that the common ancestors of GBNPP mountain goats likely originated from this area before splitting off and colonizing GBNPP after the Last Glacial Maximum. The grouping of outer coast Table Mountain subpopulation with two samples from Yakutat indicate that mountain goats in that area could also have a route north of the park boundary that followed the coast down to Table Mountain. The demographic analysis indicated that GBNPP was likely colonized by mountain goats between the retreat of the Last Glacial Maximum and the early stages of the Holocene. Consistent with this is data suggesting many mammalian species colonized northern Southeast Alaska during this period, including rodents (Runck and Cook 2005), mustelids (Cook et al. 2001), carnivores (Klein 1965), and ungulates (Klein 1965, MacDonald and Cook 2009). During this time, glaciers still filled many of today’s fjords, allowing for the possibility of a more direct route of colonization across the landscape.

Mountain goats in GBNPP exhibit some subtle patterns of IBD and spatial autocorrelation, which has been shown with other mountain goat populations, (Shafer et al. 2011, Parks et al. 2015, White et al. 2021a). The IBD patterns and lack of a PC versus latitude and longitude relationship indicates a relatively rapid wave of colonization across GBNPP, which is not unexpected at a small scale. Mountain goats exhibit female philopatry which results in patterns of closely related individuals staying together (Côté and Festa-Bianchet 2008). These patterns along with low *N*_*e*_ values suggest that genetic drift is the main driver of genetic diversity for these mountain goat subpopulations. Mountain goats are alpine specialists and have the unique ability to traverse across steep, rocky ‘escape terrain’ that most predators cannot access (Côté and Festa-Bianchet 2008, Shafer et al. 2012). Beyond the mountainous and forested landscapes, GBNPP is also made up of numerous deep inlets and fjords which likely act as barriers to contemporary mountain goat movement. Similar to what has been found for brown and black bears (Lewis et al. 2015, 2020), the recently deglaciated Glacier Bay fjord isolates subpopulations unless there is suitable habitat to facilitate movement around the fjord.

On the west side of Glacier Bay, mountain goats are consistently observed on Marble Mountain, despite its isolated geographic position relative to other groups of mountain goats (Lewis and White 2015). Marble Mountain had the lowest estimated genetic diversity and smallest estimated population size (n = 49; Lewis and White 2015); with few neighboring mountains inhabited by mountain goats, the amount of gene flow is likely decreased causing this reduction in genetic diversity (e.g. Shirk et al. 2010). Additionally, gene flow is not only a function of distance, but also population size, further causing small subpopulations, like that of Marble Mountain, to experience low gene flow and genetic diversity (Frankham 1996).

Interestingly, clustering analysis showed that two of the study areas (Mt. Wright and Tidal Inlet) are not genetically distinct from one another, indicating that gene flow between the study areas is either currently ongoing, separation has occurred recently, or that colonization was recent. A 2 km wide fjord separates these two areas but was covered in ice in the late 1800s and movement between them was likely more feasible during that time.

### 4.2 Response to climate change

Mountain goat populations are projected to be negatively affected by climate change (White et al. 2011, 2018) and our demographic simulation analyses suggest the same pattern is likely to occur in GBNPP. These analyses are not intended to be literal space and time predictions, but rather to heuristically examine whether predicted climate change is likely to be result in favorable or unfavorable outcomes for mountain goats in a particular area. In GBNPP, the projected decrease in winter snowfall and increase in summer temperature seen in climate models suggested climate change is likely to negatively affect mountain goats in all four study areas. These results are similar to that of White et al. (2018), who also found small populations were more strongly influenced by climate-mediated perturbations. To avoid these negative outcomes, mountain goats must either adapt to this changing environment by adjusting daily activity budgets or move to more suitable habitat, which would likely mean moving higher in elevation (Moritz et al. 2008). While behavioral adaptations are an important mechanism that can enable mountain goats to adapt to a changing climate, we have limited understanding of their ability to adapt or behaviorally mitigate these changes. The ability to move to more suitable habitat becomes increasingly important as less habitat is available. Unfortunately, the cost of travel across the landscape will likely increase because of climate change (Figure 3), further highlighting the negative impact that climate change will have on mountain goats.

Surprisingly, genetic variation was correlated more with Euclidean distance than LCP distance (Table S3). The LCP is reflective of the contemporary landscape and climatic patterns, which often correlates better than geographic distance in mountain goats (Shirk et al. 2010, Shafer et al. 2012). This correlation between genetic variation and Euclidean distance rather than LCP distance variation supports our assertation that historical patterns of colonization are the primary drivers of contemporary genetic patterns. During past glaciation events, ice filled the fjords, allowing for a more direct route of colonization: this is reflected in the Euclidean distances, all of which encompass the fjords when comparing the Mt. Wright to Marble Mountain subpopulations, for example. Since the ice has retreated, the landscape and movement corridors have changed and main fjord which allowed for the direct route is no longer traversable, thus explaining the lack of correlation between contemporary LCP and genetic variation. Any contemporary and future movement, however, will rely on the current landscape and corridors (e.g. Shafer et al. 2012, Wolf et al. 2020); thus we would expect the future LCP analysis to be more important for predicting future mountain goat connectivity. Additional research on shifts in mountain goat movement across the landscape would be valuable for further understanding how climate change will impact populations in Southeast Alaska. One limitation to note is that locations were used from samples collected during late winter-summer which potentially missed the full range of mountain goat habitat. For future studies on climate-induced changes in mountain goat habitat, we suggest collecting representative year-round samples and designing a complex model that incorporates finer-scale, climate-sensitive behavioral strategies to thermal stress, explicitly considers seasonal differences in ecology and distribution, and considers local climates and associated trajectories. Nevertheless, we feel that this model does provide valuable insight into future climatic changes that will affect mountain goat survival and movement across the landscape.

Movement corridors are crucial for maintaining connectivity and gene flow for isolated subpopulations on patchy landscapes (Taylor et al. 1993, Kahilainen et al. 2014, Schlaepfer et al. 2018). Connectivity across a landscape can be obstructed by the addition of roads or development which have been found to hinder mountain goat movement (Shirk et al. 2010). The construction of new trails or increased tourism could also affect the landscape connectivity for mountain goats and should be considered when making management decisions. Ecological niche modeling suggested that the climate envelope mountain goats currently use in this area will be greatly reduced and shifted, and may increase the cost of travel (i.e. connectivity) will increase. With the incorporation of genetic information to the current knowledge of population dynamics of mountain goats on a small scale, land managers can make more informed decisions to minimize the disturbance on subpopulations of mountain goats that are more vulnerable to disturbance.

## CRediT authorship contribution statement

**Kiana B. Young:** Conceptualization, Methodology, Software, Validation, Formal analysis, Investigation, Resources, Data Curation, Writing – Original Draft, Writing – Review & Editing, Visualization, Project administration. **Tania M. Lewis:** Conceptualization, Writing – Review & Editing, Supervision, Project administration, Funding acquisition. **Kevin S. White:** Methodology, Writing – Review & Editing. **Aaron B. A. Shafer:** Conceptualization, Methodology, Resources, Writing – Review & Editing, Supervision, Project administration, Funding acquisition.

## Acknowledgements

This work was funded by the CFI-JELF (36905; A.B.A.S.), Compute Canada Resources for Research Groups (GME-665-01; A.B.A.S.), NSERC Discovery Grant (A.B.A.S.: RGPIN-2017-03934) and Ontario Early Researcher Award (A.B.A.S.: ER18-14-209). This research was conducted in collaboration with Trent University, the National Park Service, and Alaska Department of Fish & Game. Thank you to the Glacier Bay National Park and Preserve boat captains for providing us with safe transport to our study areas. We are also deeply grateful for the citizen scientists for their assistance in collecting samples in areas that we were not able to access.

