## Supplemental Information for "Quantifying the effects of recent glacial history and future climate change on a unique population of mountain goats"

**SUPPLEMENTAL TABLES**

Table S1. Proportion of individuals (n = 137) within each sampling area assigned

to each subpopulation (across the top) and proportion of individuals in each

subpopulation that are admixed (q < 0.8) across 8 sampling areas in GBNPP and surrounding areas.

| *Sampling Area* | *K1* | *K2* | *K3* | *K4* | *K5* |
| --- | --- | --- | --- | --- | --- |
| Yakutat | 0.00 | 0.00 | 0.00 | 1.00 | 0.00 |
| Chilkat | 0.83 | 0.00 | 0.17 | 0.00 | 0.00 |
| GLB | 0.32 | 0.04 | 0.52 | 0.02 | 0.10 |
| Marble Mtn | 0.08 | 0.92 | 0.00 | 0.00 | 0.00 |
| Mt. Wright | 0.30 | 0.00 | 0.10 | 0.00 | 0.60 |
| Muir | 0.00 | 0.00 | 1.00 | 0.00 | 0.00 |
| Table Mtn | 0.00 | 0.00 | 0.00 | 1.00 | 0.00 |
| Tidal Inlet | 0.05 | 0.05 | 0.00 | 0.10 | 0.80 |
| Admixed | 0.41 | 0.16 | 0.38 | 0.26 | 0.33 |

Table S2. Environmental variables and their description, percent contribution, and permutation importance used in Maxent analysis. Only those variables marked with an asterisk (*) were retained in the analysis.

| *Variable* | *Description* | *Percent contribution* | *Permutation importance* |
| --- | --- | --- | --- |
| BIO1* | Annual Mean Temperature | 3.5 | 13.0 |
| BIO2* | Mean Diurnal Range (Mean of monthly (max temp - min temp)) | 3.3 | 10.0 |
| BIO3* | Isothermality (BIO2/BIO7) (×100) | 10.7 | 0.1 |
| BIO4* | Temperature Seasonality (standard deviation ×100) | 2.4 | 8.6 |
| BIO5 | Max Temperature of Warmest Month | - | - |
| BIO6 | Min Temperature of Coldest Month | - | - |
| BIO7 | Temperature Annual Range (BIO5-BIO6) | - | - |
| BIO8* | Mean Temperature of Wettest Quarter | 4.4 | 0.4 |
| BIO9* | Mean Temperature of Driest Quarter | 7.0 | 11.8 |
| BIO10* | Mean Temperature of Warmest Quarter | 2.3 | 8.2 |
| BIO11* | Mean Temperature of Coldest Quarter | 9.5 | 5.2 |
| BIO12 | Annual Precipitation | - | - |
| BIO13* | Precipitation of Wettest Month | 2.7 | 6.0 |
| BIO14* | Precipitation of Driest Month | 9.6 | 24.2 |
| BIO15* | Precipitation Seasonality (Coefficient of Variation) | 14.1 | 4.3 |
| BIO16 | Precipitation of Wettest Quarter | - | - |
| BIO17 | Precipitation of Driest Quarter | - | - |
| BIO18 | Precipitation of Warmest Quarter | - | - |
| BIO19* | Precipitation of Coldest Quarter | 4.0 | 4.0 |
| Heat Load Index* | Measure of incident radiation based off slope and aspect | 0.7 | 0.1 |
| Landcover* | Vegetation type and landcover | 25.4 | 2.8 |

Table S3. Models tested for multiple regression on distance matrices. ‘Genetic’ indicates the genetic distance matrix using Moran’s *I*. ‘Euclidean’ indicates the Euclidean distance between points. ‘LCP’ indicates the current least cost path value matrix.

| *Model* | *R^2^* | *P* |
| --- | --- | --- |
| genetic ~ euclidean | 0.217 | 0.001 |
| genetic ~ LCP | 0.001 | 0.001 |
| euclidean ~ LCP | 0.002 | 0.113 |
| genetic ~ LCP + euclidean | 0.217 | 0.001 |

**SUPPLEMENTAL FIGURES**

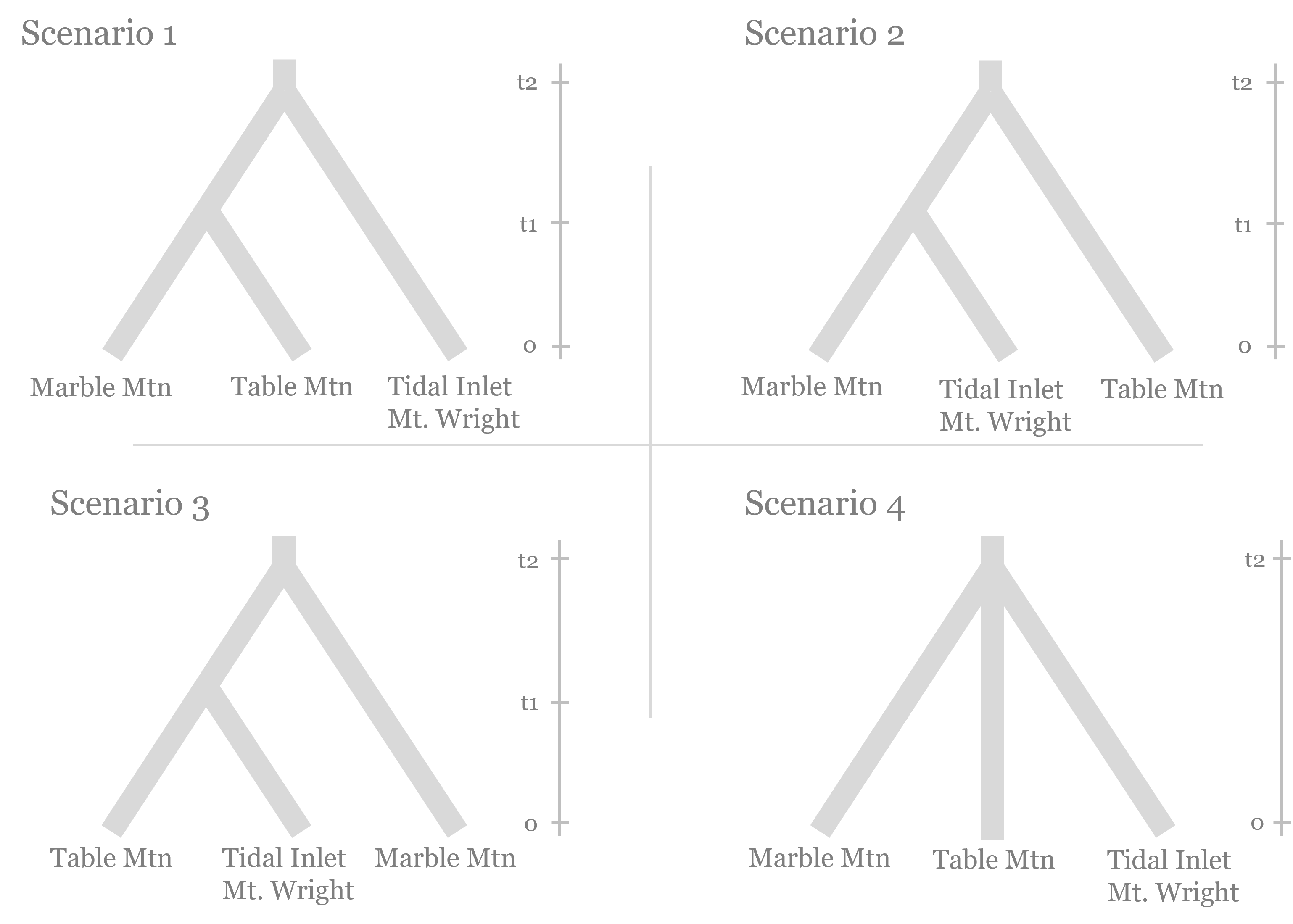

Figure S1. Scenarios tested for demographic analysis of subpopulations using computation software DIYABC (v2.0, Cornuet et al. 2014) to reconstruct the demographic history within Glacier Bay National Park and Preserve, Alaska.

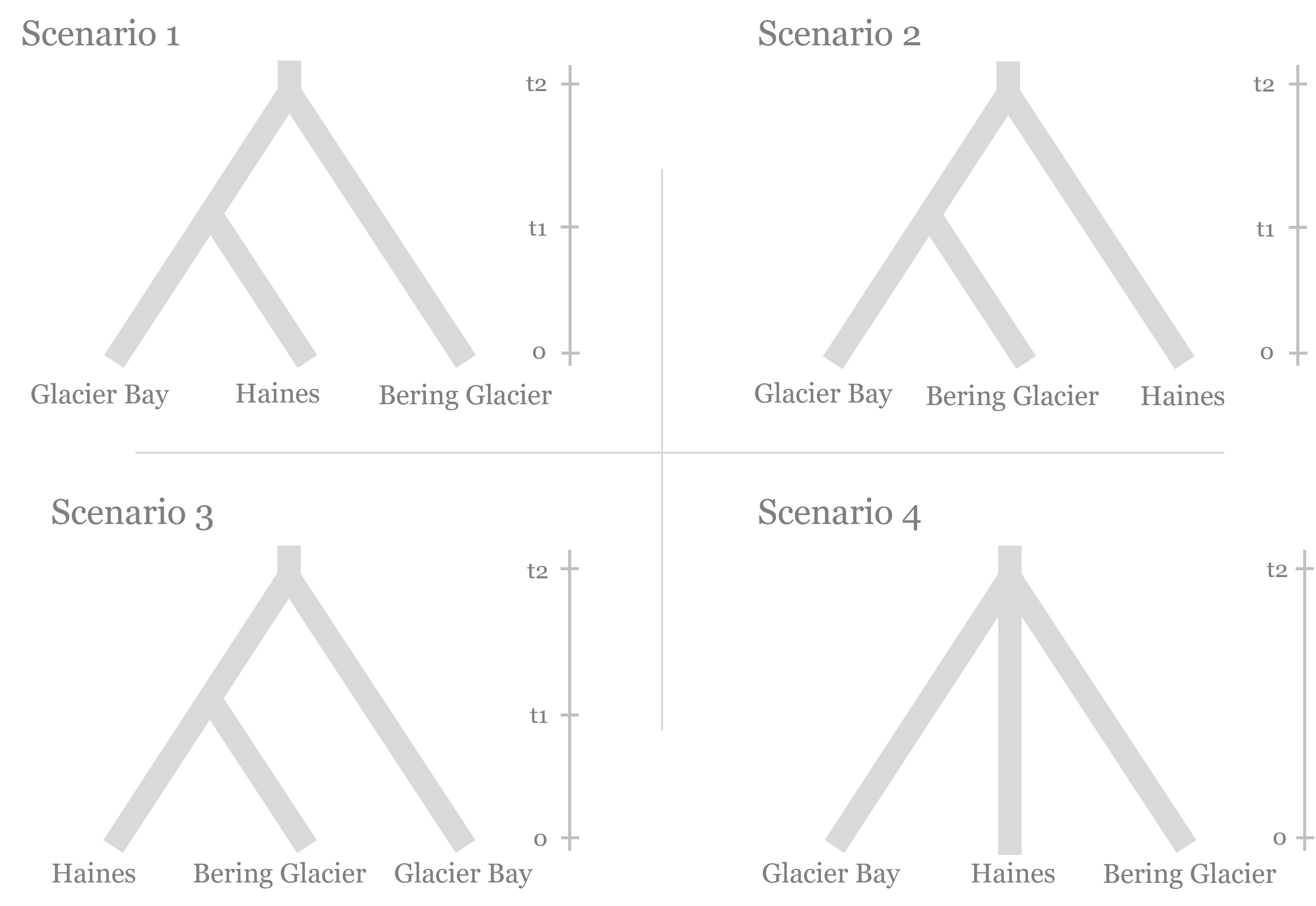

Figure S2. Scenarios tested for demographic analysis of Glacier Bay National Park and Preserve, Alaska and surrounding areas using computation software DIYABC (v2.0, Cornuet et al. 2014).

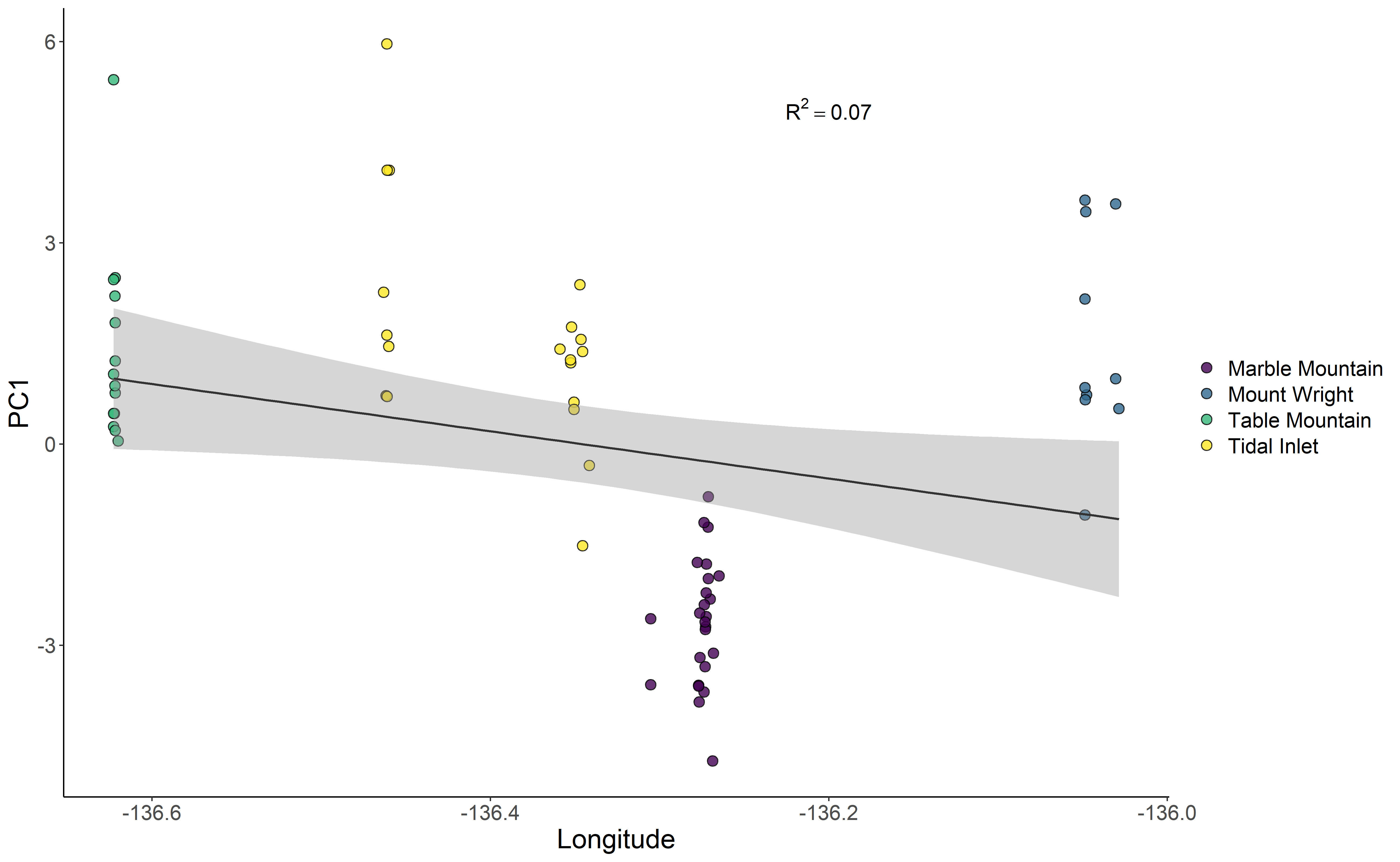

Figure S3. PC1 versus longitude for mountain goat allele frequencies (n = 68) for the four study areas identified in Glacier Bay National Park and Preserve, Alaska (indicated by color).

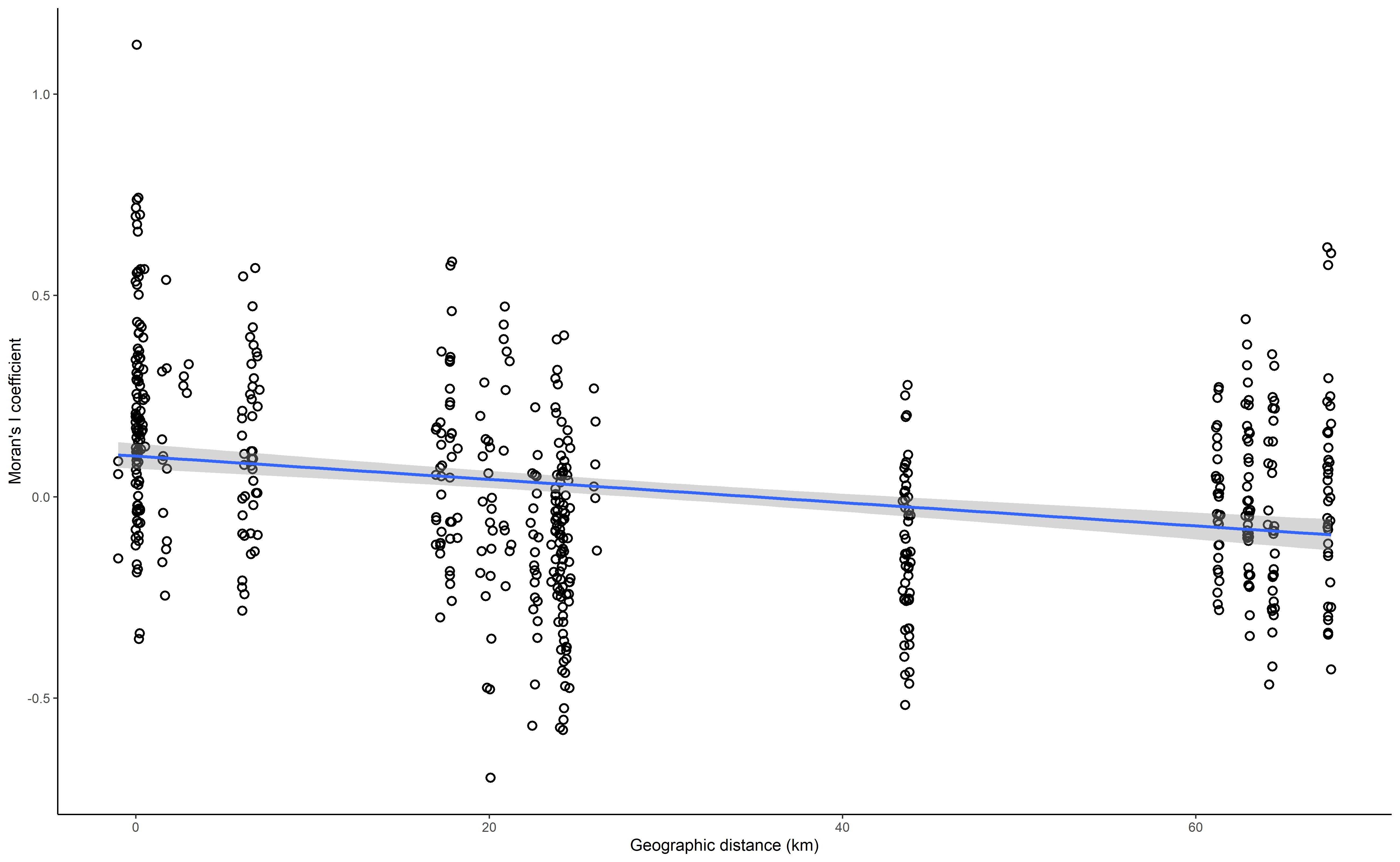

Figure S4. Isolation-by-distance as measured by Moran’s *I* of pairwise genetic relatedness versus the Euclidean geographic distance in kilometers.

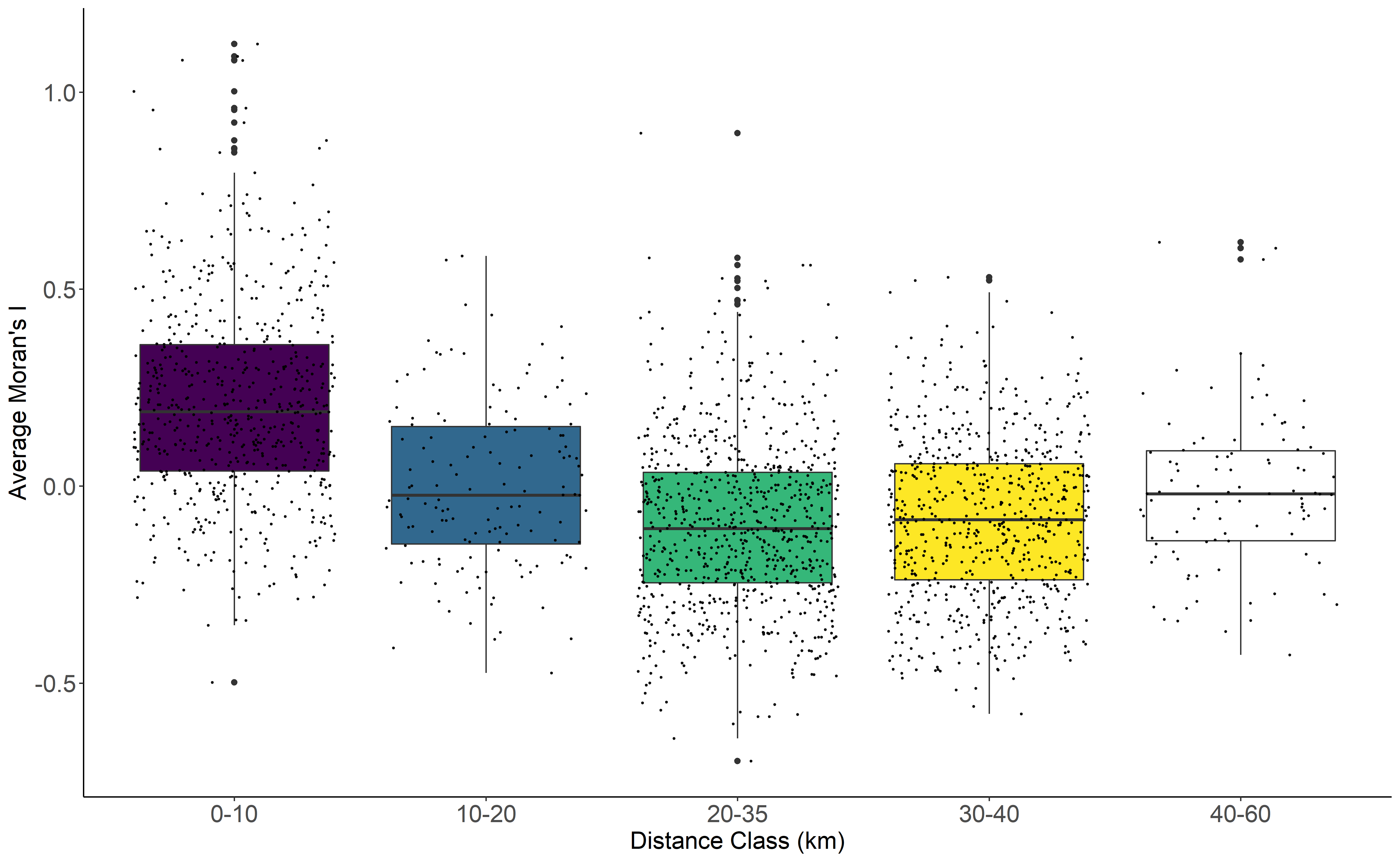

Figure S5. Spatial autocorrelation for mountain goats from four study areas in Glacier Bay National Park and Preserve, Alaska (n = 68). Moran’s *I* data generated from the program SPAGeDi.

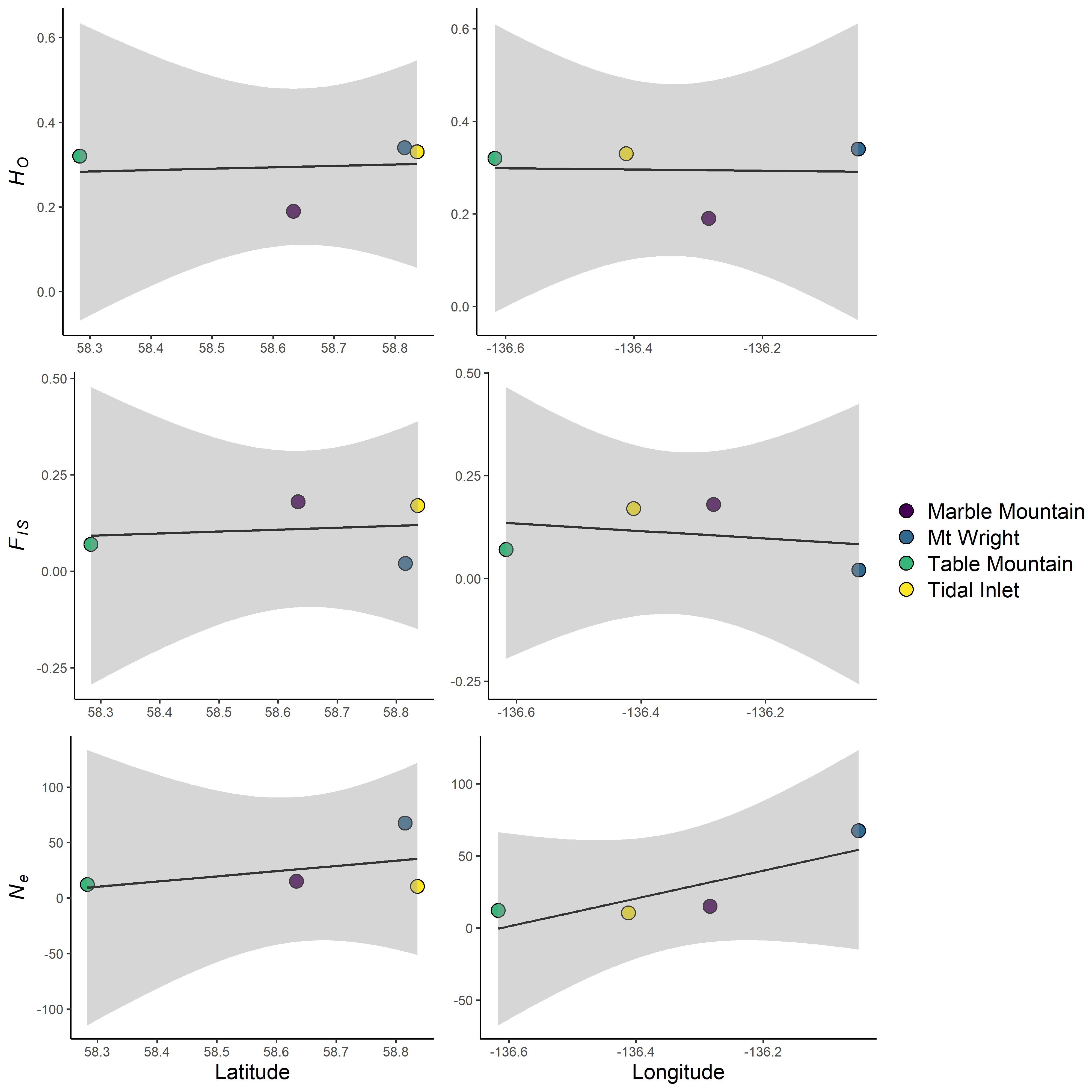

Figure S6. Latitude and longitude plotted against *F_IS_*, *H_O_*, and *N_e_* for mountain goat subpopulations in Glacier Bay National Park, Alaska. Sampling areas are indicated by color.

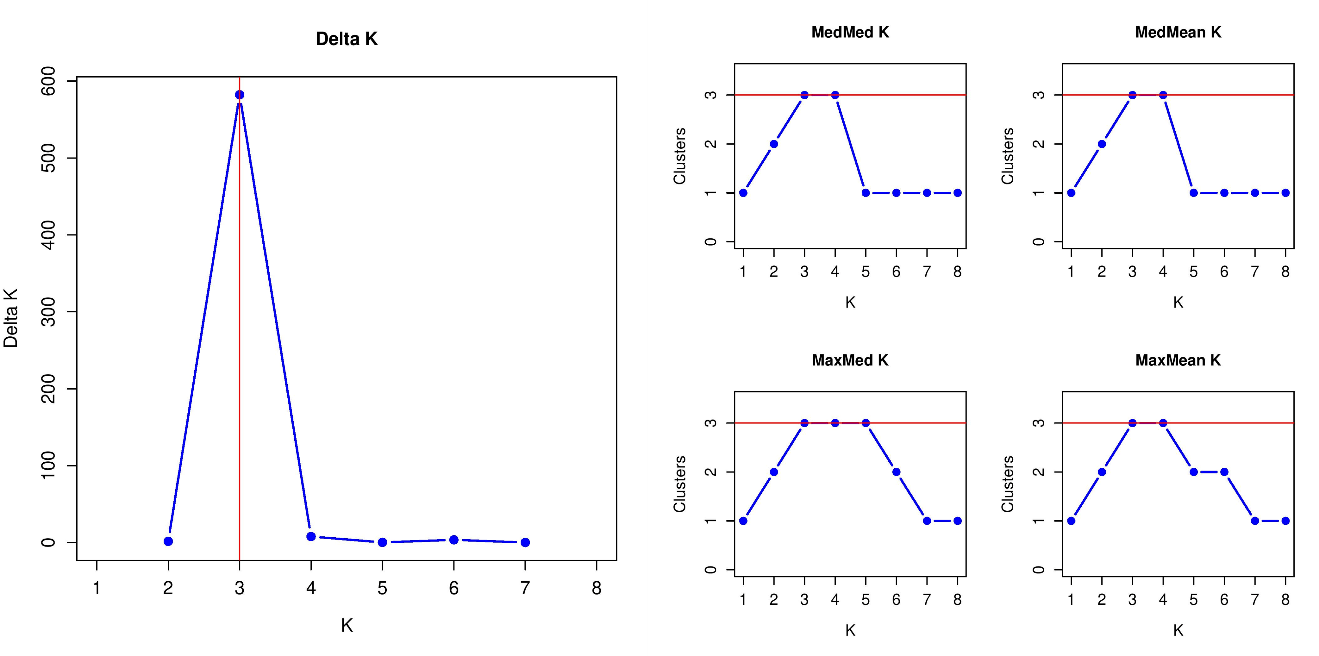

Figure S7. The number of subpopulations of mountain goats in Glacier Bay National Park and Preserve, Alaska according to the Evanno et al. (2005) and Puechmaille (2016) methods.

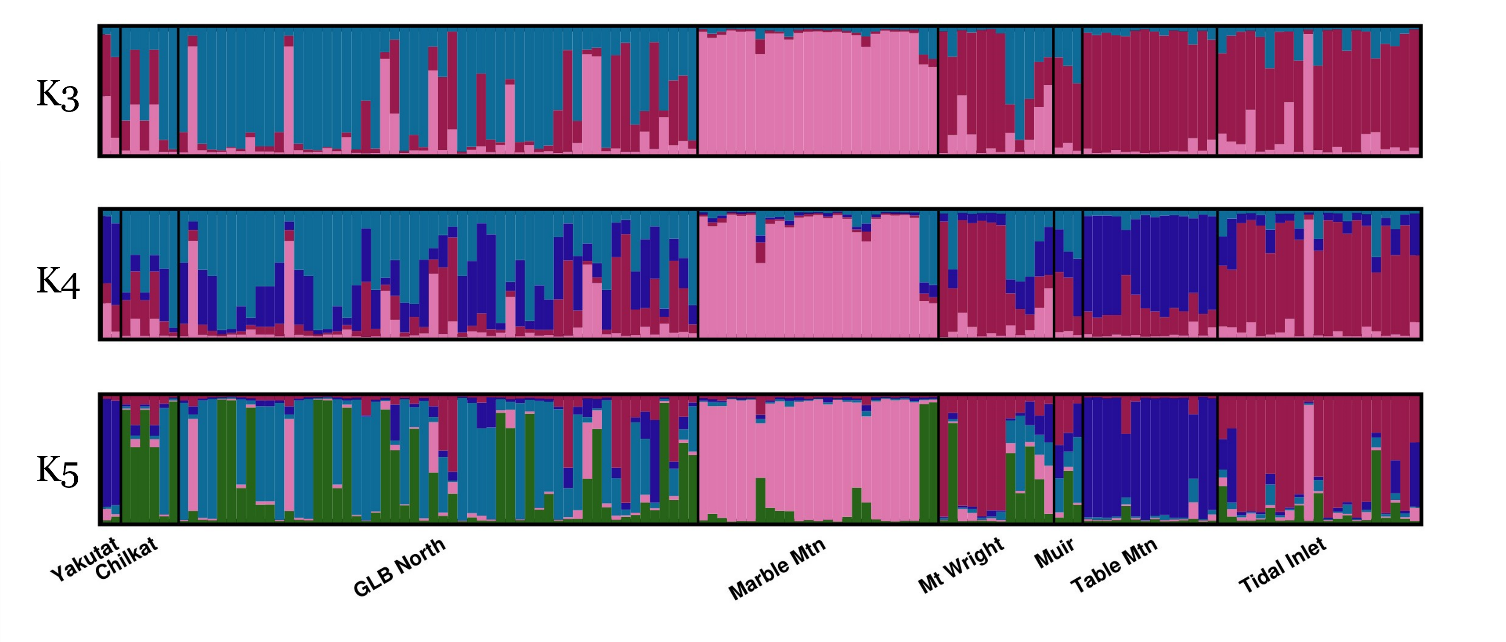

Figure S8. STRUCTURE plots for K = 2, K = 3, and K = 4 for mountain goat samples (n = 137) across eight locations in and around Glacier Bay National Park and Preserve, Alaska. Marble Mtn. Mt.Wright, Table Mtn, and Tidal Inlet are the four focal study areas within Glacier Bay National Park and Preserve.
